# Relationship between Students’ Attitude towards, and Performance in Mathematics Word Problems

**DOI:** 10.1101/2022.11.21.517411

**Authors:** Robert Wakhata, Sudi Balimuttajjo, Védaste Mutarutinya

**Affiliations:** African Centre of Excellence for Innovative Teaching and Learning Mathematics and Science (ACEITLMS). University of Rwanda, College of Education (UR-CE), P.o Box 55 Rwamagana, Rwanda, https://ur.ac.rw; Mbarara University of Science and Technology; P.o Box 1410 Mbarara, Uganda. https://www.must.ac.ug; University of Rwanda, College of Education (UR-CE), P.o Box 55 Rwamagana, Rwanda. https://ur.ac.rw

**Keywords:** attitude, mathematics, linear programming, secondary schools, performance

## Abstract

The study explored the direct and indirect relationship between students’ attitude towards, and performance in mathematics word problems (MWTs), mediated by the active learning heuristic problem solving (ALHPS) approach. Specifically, this study investigated the correlation between students’ performance and their attitude towards linear programming (LP) linear programming word tasks (ATLPWTs). Tools for data collection were: the adapted Attitude towards Mathematics Inventory-Short Form (ATMI-SF), (α=.75) as a multidimensional measurement tool, and linear programming achievement tests (pre-test and post-test). A quantitative approach with a quasi-experimental pre-test, post-test, and non-equivalent control group study design was adopted. A sample of 60811th-grade grade Ugandan students (291 male and 317 female) from eight secondary schools (both public and private) participated. Data were analyzed using PROCESS macro (v.4) for SPSS version 26. The results revealed a direct significant positive relationship between students’ performance and their ATLPWTs. Thus, it is likely that students’ attitude positively and directly impacted their performance. The present study contributes to the literature on performance and attitude towards learning mathematics. Overall, the findings carry useful practical implications that can support the theoretical framework for enhancing students’ performance and attitude towards mathematics word problems.

## 1.0 Introduction and the Concept of Students’ Attitude towards Mathematics

The term attitude is the most indispensable concept in contemporary social psychology and science. It is related to emotional and mental entities that drive an individual towards performing a particular task (Perloff, 2016). According to Aiken (1970) attitude is “a learned disposition or tendency on the part of an individual to respond positively or negatively to some object, situation, concept or another person” (p. 551). Lin and Huang (2014), define attitude towards mathematics as positive, negative, or neutral feelings and dispositions. Attitude can be bi-dimensional, (a person’s emotions and beliefs) or multidimensional (affect, behavior, and cognition). Over the last decades, an extensive body of research from different settings and contexts has investigated variables that influence students’ attitude towards Science, Technology, Engineering and, Mathematics (STEM) (e.g., Aiken 1970; Gardner, 1975; Kempa & McGough, 1977). This means that attitude determines and may be used as a predictor for academic achievement. In this study, we are particularly concerned with an exploration of the effect of the heuristic problem-solving approach on students’ attitude towards learning mathematics, and the topic of LP in particular. This is due to significant roles LP play in constructing elementary and advanced models for understanding science, technology and engineering (STE).

Numerous studies on students’ attitude towards mathematics which is always translated as liking and disliking of the subject have been published (e.g., Arslan et al., 2014; Davadas & Lay, 2020; Pepin, 2011; Utsumi & Mendes, 2000). To some secondary school students, mathematics appears abstract, difficult to comprehend, boring and viewed with limited relationship or relevance to everyday life experiences. Students start learning the subject well but gradually start disliking some topics or the entire subject. They feel uncomfortable and nervous during learning and examinations. This is partly attributed to students’ lack of self-confidence, and motivation during problem-solving. To some students, persevering and studying advanced mathematics has become a nightmare. Indeed, some students do not seem to know the significance of learning mathematics beyond the compulsory level. Students may (may not) relate mathematical concepts beyond the classroom environment if they have a negative attitude towards the subject. This may lead to their failure to positively transfer mathematical knowledge and skills in solving societal problems.

Mathematicians have attempted to research and understand affective variables that significantly influence students’ attitude towards mathematics(e.g., Barmby et al., 2008; Davadas & Lay, 2020; Di Martino & Zan, 2011; Evans & Field, 2020; Grootenboer & Hemmings, 2007; Hannula, 2002; Maamin et al., 2022; Marchis, 2011; Pongsakdi et al., 2019; Yasar, 2016; Zan et al., 2006). Some researchers have gone ahead to ask fundamental questions on whether or not students’ attitude towards mathematics is a general phenomenon or dependent on some specific variables. To this effect, some empirical findings report students’ attitudes towards specific units or topics in mathematics to enhance the learning of that specific content and mathematics generally(e.g., Arslan et al., 2014; Estrada & Batanero, 2019; Gagatsis & Kyriakides, 2000; Julius et al., 2018; Mumcu & Aktaş, 2015; Selkirk, 1975; Townsend & Wilton, 2003).

Rather than investigating students’ general attitudes toward mathematics, recent research has also attempted to identify background factors that may provide a basis for understanding students’ attitude towards and performance in learning mathematics. Thus, students at different academic levels may have negative or positive attitude towards mathematics due to fundamentally different reasons. Yet, some empirical studies have shown existence of significant relationships between students’ attitude and performance in mathematics (e.g., Berger et al., 2020; Chen et al., 2018; Davadas & Lay, 2020; Grootenboer & Hemmings, 2007; Hwang & Son, 2021; Lipnevich et al., 2011a; Ma, 1997; Maamin et al., 2022; Mazana et al., 2018b; Mulhern & Rae, 1998; Opolot-okurut, 2010; Sandman, 1980; Tapia, 1996). From the above studies, it appears that multiple factors ranging from students’ demographic factors to teachers’ classroom instructional practices influence students’ attitude towards learning mathematics.

This paper presents results from a more specific investigation into the relationship between students’ attitude towards, and performance in linear programming mathematics word tasks (Appendix 1). This is because studies concerning attitudes towards and achievement in mathematics have begun to drift from examining general mathematics attitudes to a more differentiated conceptualization of specific students’ attitude formations, and in different units (topics). Although different attitudinal scales (e.g., Code et al., 2016; Fennema & Sherman, 1976; Tapia, 1996) were developed to measure different variables influencing students’ attitude towards mathematics, this study investigated the influence of some of these constructs on students’ performance in linear programming. According to the above authors (and other empirical findings), students’ attitude towards mathematics may be the consequence of general and specific latent factors. Thus, attitude towards learning LP word tasks was investigated with specific reference to students’ performance, controlling for other variables e.g., learning strategies, students’ gender, school type, and school location.

### 1.1 Mathematics Word Problems

Verschaffel et al., 2010) define word problems as “verbal descriptions of problem situations wherein one or more questions have raised the answer to which can be obtained by the application of mathematical operations to numerical data available in the problem statement.” The authors categorized word problems based on their inclusion in real-life world scenarios. Thus, mathematics word problems play significant roles in equipping learners with the basic knowledge, skills, and, understanding of problem-solving and mathematical modeling. Some empirical findings (e.g., Boonen et al., 2016) show that mathematics word problems link school mathematics to real-life world applications. However, the learning of mathematics word problems and related algebraic concepts is greatly affected by students’ cognitive and affective factors (Awofala, 2014; Jupri & Drijvers, 2016; Pongsakdi et al., 2020). Mathematics word problems are an area where the majority of students experience learning obstacles (Abdullah et al., 2014; Awofala, 2014; Dooren et al., 2018; Goulet-Lyle et al., 2020; Julius et al., 2018a; Pearce et al., 2011; Sa’ad et al., 2014; Verschaffel et al., 2010, 2020a, 2020b). By contrast, comprehension of mathematics word problems mainly accounts for students’ difficulties. Consequently, this has undermined students’ competence, confidence and achievement in word problems and mathematization in general.

Yet, mathematical word problems are intended to help learners to apply mathematics beyond the classroom in solving real-life-world problems. Verschaffel et al. (2020) and Boonen et al. (2016) have argued that mathematics word problems are difficult, complex, and pause comprehension challenges to most learners. This is because word problems require learners to understand and adequately apply previously learned basic algebraic concepts, principles, rules and/or techniques. Indeed, most learners find it difficult to understand the text in the word problems before transformation into models. This is partly due to variation in their comprehension abilities and language (Strohmaier et al., 2020). Consequently, learners fail to write required mathematical algebraic symbolic operations and models. Yet, incorrect models lead to wrong algebraic manipulations and consequently wrong graphical representations and solutions.

Notably, research findings by Meara et al. (2019), and Evans and Field (2020) indicate that students’ mathematical proficiency is partly explained by their transitional epistemological and ontological challenges from primary to secondary education. Other studies (e.g., Georgiou et al., 2007; Grootenboer & Hemmings, 2007; Li et al., 2018; Norton, 1998; Sherman, 1979; Sherman, 1980) attribute students’ performance and achievement in mathematics to gender differences. Research by (Luneta, 2022) shows that the challenges experienced in mathematics education are a by-product of those in education in general, and they span from policy, curriculum, instruction, learning, and information technology to infrastructure. Thus, students may start excellently learning and performing mathematics from primary but gradually lose interest in some specific units and finally in mathematics generally. Several strategies have been adopted and/or adapted to boost students’ attitude in specific topics. For the case of LP, it is likely that students’ attitude towards mathematics and in equations, inequalities and LP in particular gradually drop in favor of other presumably simpler topics. To boost performance in mathematics word problems, Goulet-Lyle et al. (2020) proposed step-by-step problem-solving strategies to enhance mastery and performance

Students’ attitudes should, therefore, be investigated as well as their influence on their conceptual changes. Several empirical studies have also investigated the relationship between attitude towards, and achievement in mathematics across all levels, and in different contexts (e.g., Bayaga & Wadesango, 2014; Camacho et al., 1998; Chun & Eric, 2011; Davadas & Lay, 2020; Karjanto, 2017; Khavenson et al., 2012; Ozdemir & Ovez, 2012; Quaye, 2015; Selkirk, 1975; Tahar et al., 2010; Utsumi & Mendes, 2000; Yáñez-Marquina & Villardón-Gallego, 2016). In particular, these studies generally focused on students’ attitude towards mathematics, and many of them from the western context (Kasimu &Imoro, 2017). Yet, students may have different perceptions and attitudes towards specific content (topics) in mathematics irrespective of their setting, context and the learning environment.

To enhance mathematical conceptual proficiency, educators should boost students’ cognitive and affective domains in specific mathematics content. This is because students’ affective domain may directly influence their cognitive and psychomotor domains. In this study, it was predicted that students’ proficiency in solving LP word tasks is largely influenced by their attitude and prior algebraic knowledge, skills, and experiences. Julius et al. (2018a), and (2018b) noted that prior conceptual understanding coupled with students’ attitudes towards solving algebraic concepts impacted students’ inherent procedures in writing relational symbolic mathematical models from word problems, and provision of correct numerical solutions. Despite numerous difficulties encountered by students in algebraic inequalities (e.g., Fernández & Molina, 2017; Molina et al., 2017;Bazzini & Tsamir, 2004; Tsamir & Almog, 2001; Tsamir & Bazzini, 2004, 2006; Tsamir & Tirosh, 2006), a combination of methods (strategies) rather than one specific method can be applied to overcome specific students’ learning challenges.

### 1.2 Attitude towards Mathematics Word Problems

Linear programming, a topic in secondary school mathematics (and also as a course unit at University) is one of the algebraic topics that require students’ understanding of basic mathematical principles and rules. It is a basic introduction to other advanced methods for solving and optimizing LP problems. Linear programming is a classical unit, “the cousin” of mathematics word problems which has gained significant applications in the last decades in mathematics, science, and technology (Aboelmagd, 2018; Colussi et al., 2013; Parlesak et al., 2016; Romeijn et al., 2006). This is because LP links theoretical to practical mathematical applications. The topic provides elementary modeling skills for later applications (Vanderbei, 2014). Previous empirical studies have revealed that LP and/or related concepts are not only difficult for learners but also challenging to teach (Awofala, 2014; Goulet-Lyle et al., 2020; Kenney et al., 2020; Verschaffel et al., 2020a, 2020b). Different factors account for learners’ challenges in mathematics word problems (Ahmad et al., 2010; Haghverdi et al., 2012; Heydari et al., 2015). The challenges range from students’ comprehension of word problem statements, their attitude towards the topic, to their transformation from conceptual to procedural knowledge and understanding. Learners’ attitudes toward solving algebraic word problems should, therefore, be investigated and integrated during classroom instruction to help educational stakeholders provide appropriate and/or specific instructional strategies.

### 1.3 The Ugandan Context

In Uganda, studies on predictors of students’ attitude towards science and mathematics are scanty. There are no recent empirical findings on students’ attitude or performance in mathematics and mathematics word problems in particular. Solving LP tasks (by graphical method) is one of the topics taught to the 11th grade Ugandan lower secondary school students (NCDC, 2008, 2018). Despite students’ general and specific learning challenges in mathematics, the objectives of learning LP are embedded within the aims of the Ugandan lower secondary school mathematics curriculum. Some of the specific aims of learning mathematics in Ugandan secondary schools include…enabling individuals to apply acquired skills and knowledge in solving community problems, instilling a positive attitude towards productive work…” (NCDC, 2018). Generally, the learning of LP word problems aims to develop students’ problem-solving abilities, application of prior algebraic conceptual knowledge and understanding of linear equations and inequalities in writing models from word problems, and from real-life-world problems. Despite the learning challenges, the topic of LP is aimed at equipping learners with adequate knowledge and skills for doing advanced mathematics courses beyond the minimum mathematical proficiency at Uganda Certificate of Education (UCE).

However, every academic year, the Uganda National Examinations Board (UNEB) highlights students’ strengths and weaknesses in previous examinations at UCE. The consistent reports (UNEB. 2016, 2018, 2019, 2020) on previous examinations on work of candidates show that students’ performance in mathematics is not satisfactory especially at distinction level. In particular, previous examiners’ reports show students’ poor performance in mathematics word problems. The examination reports further revealed numerous students’ specific deficiencies in the topic of LP (Appendix1). Students’ challenges in LP mainly stem from comprehension of word problems to formation of wrong linear equations and inequalities (in two dimensions) from the given word problem in real-life situations. Thus, wrong models derived from questions may result in incorrect graphical representations, and consequently wrong solutions and interpretations of optimal solutions. These challenges (and others) may consequently hinder and/or interfere with students’ construction of relevant models in science, mathematics and technology. Moreover, learners have consistently demonstrated cognitive obstacles in answering questions on LP, while others elude these questions (appendix 1) during national examinations. Noticeably absent in all the UNEB reports are factors that account for students’ weaknesses in learning LP and the specific interventions to overcome students’ challenges in LP. Some students have developed a negative attitude towards the topic. Yet, students’ attitudes may directly impact on their learning outcomes (Code et al., 2016).

Several attitudinal scales (with both cognitive and behavioral components) have been developed (Lim & Chapman, 2013; Yáñez-Marquina & Villardón-Gallego, 2016)adopted or adapted(Lin & Huang, 2014) to assess students’ attitude towards mathematics and in specific mathematics content. For instance, Geometry Attitude Scales (Avcu & Avcu, 2015), Statistics Attitude Scales (Ayebo et al., 2019; Khavenson et al., 2012), Attitudes toward Mathematics Word Problem Inventory(Awofala, 2014), the Attitude Towards Geometry Inventory (ATGI) instrument(Utley, 2007), and others. In this study, we adapted the Attitude towards Mathematics Inventory-Short Form (ATMI-SF) instrument (Lin & Huang, 2014) to investigate the effect of the heuristic problem-solving approach on students’ attitude towards learning LP word problems. Specifically, this research explored the 11th-grade students’ attitude towards learning LP word problems (see appendix1). Taken together, research shows that a high percentage of educational stakeholders around the world are concerned about performance and attitude towards mathematics word tasks in particular. However, to fully understand students’ attitude towards and performance in mathematics, it is necessary to investigate beyond general mathematics attitudes and examine specific underlying aspects for these attitudes.

### 1.4. Objectives of the Study

The purpose of this study was to explore the relationship between students’ attitude towards, and performance in LP word problems. Specifically, the present study aimed at:

-Investigating the direct effect of students’ attitude towards, and performance in mathematics word problems.
-Exploring the indirect effect of students’ attitude towards, and performance in LP word tasks mediated by active learning heuristic problem solving strategies.

## 2.0 The Theoretical Framework

This study is situated on the theoretical framework according to constructivism, and Eccles, Wigfield, and colleagues’ expectancy-value model of achievement motivation (Wigfield, 1994; Wigfield & Eccles, 2000). The expectancy-value model is based on the expectancy-value theories of achievement. Thus, the theory is based on the premise that success on specific tasks and the values inherent in those tasks is positively correlated with achievement, and consequently students’ abilities. In the context of the attitude towards mathematics inventory-short form (ATMI-SF), the theory combines motivation, enjoyment, confidence, and usefulness, and related latent variables to explain students’ success in learning mathematics. Constructivism is a form of discovery learning which is based on the premise that teachers facilitate learning by actively involving learners so that they construct their world knowledge and understanding based on individual prior experiences and schema (Olusegun, 2015; Ültanır, 2012). This means that previous knowledge, understanding and reflection with new knowledge is inevitable for subsequent learning and acquisition of both conceptual and procedural knowledge.

We are particularly concerned about students’ latent efforts, and persistence, their perceived difficulties in learning LP word tasks and experiences learners encounter when solving LP word tasks. Students’ challenges in LP may largely depend on their insufficient previous algebraic knowledge and experiences in applying concepts of equations and inequalities. In this article we mainly discuss students’ ATLPWTs using the expectancy-value model theory within the constructivism paradigm. Using this paradigm helped to explain the ATMI-SF constructs, and their significance in enhancing the learning of mathematics in secondary schools. The expectancy-value theory and constructivism have been widely applied to enhance the learning of mathematics and science (Awofala, 2014; Fielding-Wells et al., 2017; Meyer et al., 2019; Wigfield & Eccles, 2000; Yurt, 2015). To foster a positive attitude, teachers should design different tasks for students based on their academic level and previous academic performance. This may help them to apply the previously acquired knowledge, understanding and experiences in subsequent learning. Stein et al. (2000) resonated that students’ proficiency and competency is determined by mathematical tasks they are given. Tasks at the lower cognitive stage (memorization level), for example, must be different from those at the highest cognitive level (doing mathematics). In the context of learning LP, students should first understand and appropriately apply the knowledge of equations and inequalities to adequately and proficiently solve non-routine mathematical problems.

### 2.1 Conceptual Framework of the Study

Based on the theoretical framework discussed above, a conceptual framework shows the direct relationship between students’ attitude towards, and performance in mathematics word problems, mediated by active learning heuristic problem solving strategies. The model hypothesizes the direct relationship among the stated variables. The model assumes the existence of a relationship between attitude and performance in post-test. There exist other indirect inherent relationships which are not part of this study. The model is shown in Figure 1 below. Active learning develops students’ higher-order thinking skills (analysis, synthesis, and evaluation) and also prepares them to apply mathematics in real-life scenarios. Several studies have shown that students in typical active learning classrooms perform better than those taught conventionally (Freeman et al., 2014; Prince, 2004). This is because they have an opportunity to reflect, conjecture, or predict outcomes, and then to share and discuss their concepts with teachers and their peers to activate and re-activate their cognitive processes. Active learning helps students to reflect on their understanding by encouraging them to make connections between prior mathematical knowledge and new concepts.

**Figure 1.**
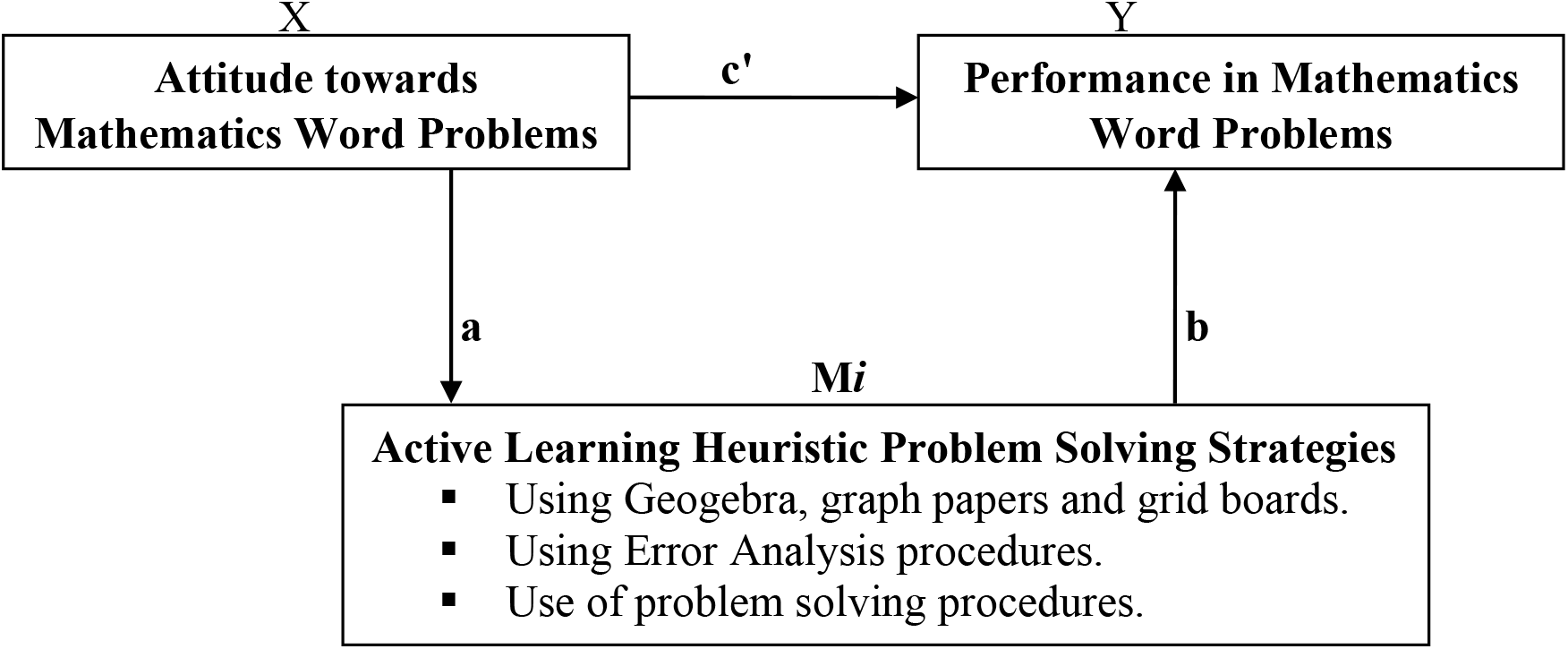
Showing Direct (**c’**) and Indirect (**b.c**) Relationship between Attitude towards and Performance in Mathematics Word Problems Mediated by Active Learning Heuristic Problem Solving Strategies

## 3.0 METHODOLOGY

A quantitative survey research design was used to collect, analyze, and describe students’ experiences and latent behavior in learning linear programming mathematics word tasks.

### 3.1 Research Design

This study investigated the relationship between students’ attitude towards, and performance in mathematics word tasks. The study adopted a quantitative approach to gain a deeper and broader understanding (Creswell, 2014; Creswell, & Plano Clark, 2018; Djamba & Neuman, 2002). A quasi-experimental pre-test, post-test, non-equivalent, non-randomized control group study design was adopted. By using the stated approach and design, researchers ably compared and contrasted students’ attitude towards learning LP word problems. Learners from the sampled schools (experimental and comparison groups), and in their intact classes participated. Intact classes were maintained to avoid interfering with the respective schools’ set timetables and schedules.

### 3.2 Sample and Participants

The sampling frame was the 11th grade students for the academic year 2020/2021. Both private and public secondary schools (rural and urban) participated. This study comprised of a research study of 608 11th grade students from eight randomly selected secondary schools. Four schools were selected from Mbale district, eastern Uganda, and four from Mukono district, central Uganda. Schools were allocated to the experimental and comparison groups by a toss of a coin. Two schools from each region were assigned to the experimental group. Selection of the 11 grade students was based on the content covered in this class as outlined in the Ugandan mathematics curriculum materials (NCDC, 2018). Of the 608 students, 291 (47.86%) were males and 492 (57.8%) were females with a mean age of 18.36 (S.D = 0.94) years. Three hundred seventeen (52.1%) students were from the comparison group while two hundred ninety-one (47.9%) were from the treatment group. The selection of students from the two distant schools within/outside the regions and assigning them to treatment group was to avoid spurious results. A situation where a particular school had more than one class “stream”, at least one hundred students with varied academic abilities were randomly selected from different classes to respond to the attitudinal questionnaires. The school (subject) heads revealed that the mathematics syllabus containing LP word problems (Appendix 1) had been completed at the time of data collection. The students were selected to provide their experiences and attitudes toward learning LP word problems. Ethical clearance was sought from research and innovation unit, at the corresponding authors’ university. Subsequent permission was sought from ministry of education, district education officers and the headteachers of sampled schools. The purpose of the study was clearly explained to the participants. Finally, identification numbers were allotted to participants before they consented. They anonymously and voluntarily completed all questionnaires.

### 3.3 Research Instruments and Procedure for administration

In addition to demographic questions, the Attitude Towards Mathematics Inventory-Short Form (ATMI-SF)(Lin & Huang, 2014), a 14-item instrument questionnaire consisting of four subscales (enjoyment, motivation, value/usefulness and self-confidence) was adapted (Appendix 2) to measure the relationship between students’ performance and their attitude towards learning LP word tasks. The ATMI-SF, a 5-point likert-type scale with response options ranging from “Strongly Disagree (1)” to “Strongly Agree (5)” was used. The ATMI-SF items were developed from (Lim & Chapman, 2013), which were also developed and validated from several mathematics attitudinal questionnaire items (Fennema & Sherman, 1976; Kasimu & Imoro, 2017; Mulhern & Rae, 1998; Primi et al., 2020; Tapia, 1996). The ATMI-SF was adapted (Appendix 2) because it directly correlates with the learning of LP, “the cousin of mathematics word problems.” English being the language of instruction in Ugandan secondary schools’ curricula, translation of questionnaire items was not required.

Content validity of the questionnaire was assessed by three experts (one senior teacher for mathematics, one senior lecturer for mathematics education, and one tutor at teacher training institution). The experts were selected based on their vast experience in teaching mathematics at various academic levels. They evaluated the appropriateness and relevance of the adapted questionnaire items. Based on their recommendations, suggestions and comments, some questionnaire items were adjusted to suit students’ academic level and language to adequately answer the research objectives. Reliability of the construct was ascertained using Cronbach alpha (0.75). This threshold is acceptable based on recommendations from Hair et al. (2010).

The ATMI-SF was administered to the treatment and comparison groups. Both the experimental and comparison groups were taught LP (equations, inequalities and LP) following the designed and approved Ugandan mathematics curriculum materials (NCDC, 2018). The experimental group was taught LP using the heuristic problem-solving approach before and after an intervention. The learning in the comparison group was purely conventional, and teachers did not follow proper and organized strategies as it was the case with the treatment group. In particular, students from the treatment group were taught LP using several active learning heuristic strategies following clearly outlined principles and strategies.

To adequately implement active learning heuristic problem-solving strategies, teachers from the treatment group were trained. First, students’ prior conceptual knowledge of equations and inequalities plus the basic algebraic principles and understanding were reviewed to link previous concepts to learning of LP. Second, several learning materials were applied to help students adequately master LP concepts. The materials included the use of graphs, grid boards, excel and GeoGebra software (see Appendix 3 for G01, G02, and G03). Empirical studies (e.g., Abdul et al., 2010; Sari et al., 2012) have found that the stated materials enhance students’ conceptual understanding and critical thinking abilities. These strategies were further integrated in problem solving strategies (Polya, 2014) by ensuring that students understand the LP word problem, devise a plan, adequately carry out the plan and finally look back to verify solution sketches (see Appendix 3 for PS01, PS02, PS03, PS04). To ensure that students minimize errors and misconceptions, the learning of LP was further integrated with Newman Error Analysis (NEA) model (Mushlihah, 2018).The instrument was designed by the researcher and validated by experts in mathematics education as an error analysis tool to provide teachers with a framework to consider the underlying reasons why students answer mathematics word problems incorrectly. The teachers emphasized question reading and decoding, comprehension, transformation, process skills and encoding (see Appendix 3 for NEA01, NEA02, NEA03, NEA04 and NEA05).

These strategies supported students’ thinking, and reasoning. Consequently, teachers’ identified students’ preconceptions, misconceptions and errors as they solved LP word tasks. Multiple representations and demonstrations were carried out to help students understand preliminary concepts and use them to optimize LP tasks. Students were engaged in the typical PS scenario individually, in pairs and in small groups. Questions and specific classroom tasks ranged from simple to complex and from concrete to abstract. Materials for instance graph papers were provided as teachers guided and demonstrated the graphical solution of LP problems. Teachers provided procedures by supporting and guiding learners during the learning process. All students’ preconceptions, misconceptions and errors were addressed. For instance, confusion on whether or not to use dotted lines or solid lines to represent the inequality y>a+bx, plotting lines to represent an equation or inequality, shading wanted or unwanted regions, etc.

All learners were assigned varied tasks to apply suitable strategies for yielding the optimal solution of a LP word problem. The “bright” students were allowed to share their experiences with the low attainers in pairs or in their small groups. Where necessary, local language was allowed for students’ peer instruction in pairs or small groups to master the underlying concepts. The classroom interaction and presentations enhanced students’ conceptual knowledge and procedural understanding of LP concepts. This further improved their reasoning, creativity and critical thinking. Consequently, constructive and informative feedback to retain their conceptual understanding was fostered. All challenging concepts on LP were harmonized by individual teachers during reflection and the evaluation phase. All students who had challenges even after this phase were allowed to repeat the tasks by doing corrections and submitting their work for remarking. The intervention took approximately three months from October 2020 to February 2021. To find out whether or not active learning heuristic problem-solving approach changed students’ attitude and performance, the post attitude questionnaire and achievement test were administered to both groups, and analysis was done by comparing and contrasting students’ feedback on attitudinal constructs and the post-test scores. Finally, this study established if there was a significant relationship between students’ performance and their attitude towards mathematics word tasks, mediated by active learning heuristic problem-solving approach (Appendix 3).

### 3.4 Data Analysis

#### Procedure

The ATMI-SF questionnaires were completed by the sampled students at their respective schools in their natural classroom setting. The questionnaires were completed in at most 30 minutes on average. The survey instrument contained a ‘filter statement’, as a Social Desirability Response (SDR) to verify and discard respondents’ questionnaires especially those who did not read (see item 15 in appendix 1) or finish answering all questionnaire items (Bäckström & Björklund, 2013; Latkin et al., 2017). Written consent was received from all participants and participation in this study was completely voluntary and confidential. Participants who felt uncomfortable to complete the questionnaire were not penalized. Data were collected with the help of mathematics heads of department who were selected from sampled schools on the basis of their expertise and experience. Participants were explained to, the purpose of the study before administering and/or filling in of questionnaire items. In the presence of the principal researcher, research assistants and some selected school administrators, participants completed and returned all the questionnaires. Descriptive and inferential statistics were used to analyze the collected data with reference to the background characteristics and stated hypothesis. Data were analyzed using the Statistical Package for Social Sciences (SPSS) version 26, with Hayes (2022) PROCESS (v.4) macro. This provided the analysis for exploring whether or not there exist a significant relationship between students’ performance and their attitude towards LP mathematics word problems.

### 3.5 Ethical Considerations

Ethical clearance was sought from the Research and Ethics committee of corresponding authors’ university. Subsequent permission was sought and granted by the permanent secretary ministry of education and sports, the district education officers, and finally from the headteachers of sampled secondary schools. Upon accessing research participants, the questionnaires were administered to the respondents during school working hours without interfering with the school set timetables. The heads of the sampled schools provided appropriate schedules and personnel to help the principal researcher together with research assistants to administer questionnaires. All participants were informed and clearly explained to the purpose of the study before participating. They were assured of confidentiality and, anonymity before they willingly consented. Those who opted not to participate even after the distribution of questionnaires were allowed to withdraw.

## 4.0 Findings and Interpretation

### Psychometric Properties

IBM SPSS (version 26) software package was used for analysis. Preliminary statistical analysis revealed no evidence of missing data due to a few cases which were ignored because they did not exceed 5% of sample cases (Barbara G. Tabachnick, 2001; Kline Rex B, 1998; Lim & Chapman, 2013). However, out of 639 questionnaires distributed, 31 questionnaires were removed because the participants did not either conform to SDR(Bäckström & Björklund, 2013; Latkin et al., 2017) or the questionnaires were incomplete. Univariate analysis was run to examine the degree of normality (Hair et al., 2010; Pallant, 2011). The indices for skewness and kurtosis were within the acceptable ranges (±2 and ±7 respectively) (Byrne, 2010; Curran et al., 1996; Hair et al., 2010). Thus, data were fairly normally distributed.

We tested the psychometric properties (reliability and factor analysis) of the two instruments. The Cronbach alpha coefficient of the adapted ATMI-SF was α=0.75. Factor analysis was performed using the principal component (with varimax rotation) (Tabachnick, 2001; Pallant, 2011; Pituch, 2016). The values obtained were consistent with Lin and Huang’s (2014), Awofala’s (2014) findings. The Kaiser-Meyer-Olkin Measure of Sampling Adequacy Test (KMO) and Bartlett’s test of sphericity was conducted. The value of KMO = 0.77> 0.60; and that of Bartlett’s test of sphericity was significant (894.349, p<0.05) indicating a substantial correlation in the data and an acceptable fit. For a self-developed standardized active learning heuristic problem-solving (ALHPS) tool, α=0.71, KMO = 0.74 > 0.60; and that of Bartlett’s test of sphericity was significant (253.092, p<0.05). Following the above recommendations, all items were found to be acceptable with adequate construct validity, internal consistency and homogeneity (Nunnally & Berstain, 1994, Pallant, 2011). Overall, these items were deemed fit to measure the relationship between active learning heuristic problem-solving and students’ attitude towards linear programming word tasks.

Table 2 and Table 3show descriptive statistics (mean, standard deviation, skewness and kurtosis). The values in the table show students’ scores on ATMI-SF and ATLPWTs. The results do not show any significant differences between the relationship between ALHPS approach and students’ attitude towards linear programming (enjoyment, motivation, usefulness and self-confidence).Indeed, both experimental and comparison groups were similar during pre-test. There was however a change in students’ ATLPWTs due to the intervention administered. The findings show that students generally held negative attitude towards learning LP word tasks. These findings are consistent with other empirical research findings (e.g., Awofala, 2014).

**Table 2.** Descriptive Statistics: Students’ ATLPWTs by Item (see Appendix 2)

| **Items** | **N** | **Mean** | **S.D** | **Skewness** | **Kurtosis** |
| --- | --- | --- | --- | --- | --- |
| A1 | 608 | 3.16 | .971 | .047 | .089 |
| A2 | 608 | 3.31 | 1.196 | -.237 | -.793 |
| A3 | 608 | 3.43 | 1.064 | -.457 | -.242 |
| A4 | 608 | 3.42 | 1.131 | -.362 | -.536 |
| A5 | 608 | 3.27 | 1.103 | -.360 | -.362 |
| A6 | 608 | 3.43 | 1.073 | -.470 | -.273 |
| A7 | 608 | 3.47 | .941 | -.582 | -.002 |
| A8 | 608 | 3.36 | 1.018 | -.463 | -.333 |
| A9 | 608 | 3.33 | .980 | -.479 | -.203 |
| A10 | 608 | 3.36 | 1.046 | -.302 | -.512 |
| A11 | 608 | 3.26 | 1.028 | -.051 | -.453 |
| A12 | 608 | 3.31 | 1.050 | -.242 | -.385 |
| A13 | 608 | 3.12 | .946 | .060 | -.302 |
| A14 | 608 | 3.29 | 1.111 | -.359 | -.577 |

**Table 3.** Descriptive Statistics: Students’ ALHPS Approach (see Appendix 3)

| **Item** | **N** | **Mean** | **S. D** | **Skewness** | **Kurtosis** |
| --- | --- | --- | --- | --- | --- |
| G01 | 608 | 3.82 | .999 | -.884 | .710 |
| G02 | 608 | 3.75 | .852 | -.451 | -.012 |
| G03 | 608 | 3.51 | 1.071 | -.482 | -.709 |
| PS01 | 608 | 2.49 | 1.205 | .776 | -.302 |
| PS02 | 608 | 2.14 | 1.013 | 1.316 | 1.881 |
| PS03 | 608 | 2.26 | .984 | 1.103 | 1.520 |
| PS04 | 608 | 2.24 | .838 | 1.354 | 3.078 |
| NEA01 | 608 | 2.28 | .919 | 1.266 | 2.225 |
| NEA02 | 608 | 2.32 | .990 | 1.164 | 1.559 |
| NEA03 | 608 | 2.42 | .912 | .846 | 1.375 |
| NEA04 | 608 | 2.31 | .999 | 1.026 | 1.238 |
| NEA05 | 608 | 2.50 | .989 | .911 | .979 |

Table 4 and Table 5 show that the active learning heuristic problem solving approach has a significant effect on attitude.

**Table 4.**
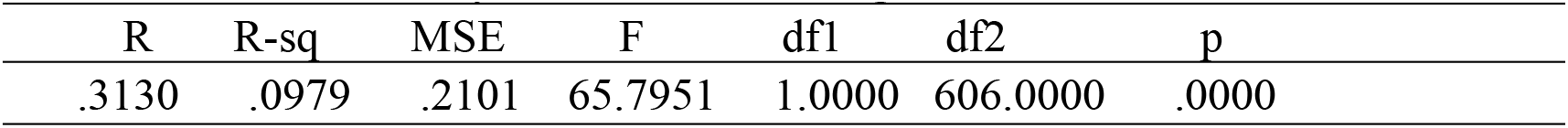
Model Summary for Active Learning.

| R | R-sq | MSE | F | df1 | df2 | p |
| --- | --- | --- | --- | --- | --- | --- |
| .3130 | .0979 | .2101 | 65.7951 | 1.0000 | 606.0000 | .0000 |

**Table 5.**
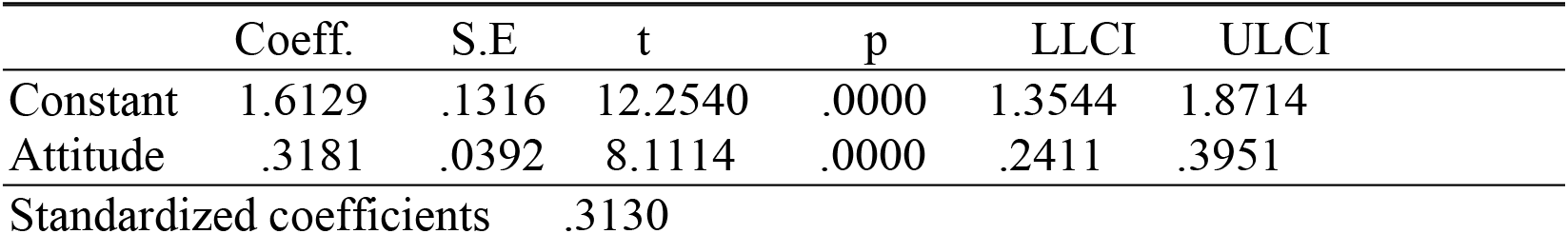
Model for Attitude towards Mathematics Word Problems.

|  | Coeff. | S.E | t | p | LLCI | ULCI |
| --- | --- | --- | --- | --- | --- | --- |
| Constant | 1.6129 | .1316 | 12.2540 | .0000 | 1.3544 | 1.8714 |
| Attitude | .3181 | .0392 | 8.1114 | .0000 | .2411 | .3951 |
| Standardized coefficients |  |  | .3130 |  |  |  |

Table 6 and Table 7 show that the attitude has a direct and significant effect on performance, and is being accounted for by approximately 50% of the model.

**Table 6.** Model Summary for Performance (Post-test)

| R | R-sq | MSE | F | df1 | df2 | p |
| --- | --- | --- | --- | --- | --- | --- |
| .7095 | .5034 | 165.0188 | 306.6195 | 2.0000 | 605.0000 | .0000 |

**Table 7.** Combined model for Active Learning and Attitude.

|  | Coeff | SE | t | p | LLCI | ULCI |
| --- | --- | --- | --- | --- | --- | --- |
| Constant | -43.9672 | 4.1202 | -10.6712 | .0000 | -52.0588 | -35.8757 |
| Attitude | 17.0392 | 1.1571 | 14.7263 | .0000 | 14.7669 | 19.3116 |
| Active Learning | 16.2792 | 1.1384 | 14.3005 | .0000 | 14.0436 | 18.5148 |

Standardized coefficients Attitude .4442, Active Learning .4314

Table 8 shows the significant direct effect of students’ attitude towards and performance in LP mathematics word tasks.

**Table 8.** Direct effect of X on Y.

| Effect | S.E | t | p | LLCI | ULCI | c' _cs |
| --- | --- | --- | --- | --- | --- | --- |
| 17.0392 | 1.1571 | 14.7263 | .0000 | 14.7669 | 19.3116 | .4442 |

However, from Table 9 and Table 10, the indirect effect of attitude on performance mediated by active learning heuristic problem-solving strategies exists with the effect size of 5.18.

**Table 9.** Indirect effect(s) of X on Y:

|  | Effect | Boot SE | Boot LLCI | Boot ULCI |
| --- | --- | --- | --- | --- |
| AL | 5.1783 | .6416 | 3.9684 | 6.4548 |

**Table 10.** completely standardized indirect effect(s) of X on Y:

|  | Effect | Boot SE | Boot LLCI | Boot ULCI |
| --- | --- | --- | --- | --- |
| AL | .1350 | .0156 | .1044 | .1659 |

## 5. Discussions and Educational Implications

This study sought to investigate the direct and indirect relationship between students’ attitude towards, and performance in LP mathematics word problems, mediated by active learning heuristic problem-solving (ALHPS) approach. This is of great educational implications to the learning of mathematics in Uganda, and supplements to other empirical findings (Opolot-okurut, 2010). Thus, altering students’ attitude may change their performance in mathematics. In this study, the psychometric properties of the adapted students’ ATLPWTs and the ALHPS approach instruments were found acceptable. The data was collected from 608 students from eight secondary schools in eastern Uganda and central Uganda. Our study revealed a direct significant positive relationship between students’ performance and their attitude towards learning mathematics word problems mediated by active learning heuristic problem solving approach.

This study indicates that students’ attitude enhanced their performance in learning LP concepts. Our findings concord with other previous empirical studies on the contribution of positive attitude in enhancing students’ performance (Chun & Eric, 2011; Juter, 2005; Lipnevich et al., 2011; Ma, 1997; Osborne et al., 2003). Seen in this way, our findings are further in agreement with the constructivism theoretical framework suggesting that educators should always review previous concepts and use them to construct new knowledge. The results of this study are likely to inform educational stakeholders in assessing students’ ATLPWTs and provide remediation and interventional strategies aimed at creating a conceptual change in students’ attitude towards LP and mathematics generally. This will further act as a lens in improving students’ achievement, as indicators of students’ confidence, motivation, usefulness, and enjoyment in learning LP word problems and mathematics generally.

These findings show that students generally had negative attitude towards LP word problems. Although some students’ ratings were below the neutral attitude (3), they indicated the usefulness of LP in daily lives. The experimental group showed a slightly favorable attitude towards LP word problems after an intervention because the problem-solving heuristic instruction was used during instruction as compared to students in comparison group who learned LP conventionally. Some students and teachers revealed that LP concepts are more stimulating, require prior conceptual knowledge and understanding of equations and inequalities, and are not interesting to learn just like other mathematics topics. The explanation provided indicated that some teachers do not adequately apply suitable instructional techniques and learning materials to fully explain the concepts. However, it was observed that teachers encouraged students to constantly practice model formation from word problems to demystify the negative belief that LP is a hard topic, thereby encouraging them to understand LP and related concepts. However, students’ performance especially those from the experimental group improved compared to their counterparts from the comparison group who almost had similar attitude towards LP before and after an intervention. The learning of mathematics should promote students’ engagement where learning is by doing through practice and that errors and misconceptions should be considered as part and parcel of learning.

The results of this study are consistent with the theoretical framework (Wigfield, 1994; Wigfield & Eccles, 2000). To achieve the purpose of this study, teachers in the experimental group varied tasks to examine students’ problem-solving and critical thinking skills. Both the experimental group and the control group acknowledged the fact that LP is a challenging topic (see appendix 1), although they highly recognized its significance in constructing models, and optimization in real life. The importance of LP rests in its application and thus teachers were tasked to help learners to develop their attitude, and conceptual understanding so that they can reason insightfully, think logically, critically and, coherently. The teachers’ competence in applying instructional strategies helped learners from the experimental group to gain deeper and broader insight, conceptual and procedural understanding, reasoning, and positive attitude. The control group in their conventional instruction still perceived LP as one of the hardest topics. Negative attitude was observed as it was indicated in the most learners’ ATMI-SF questionnaires. Thus, teachers recognized that hard work and application of prior conceptual knowledge and understanding may favorably help students to develop a positive attitude and perform better in LP. Generally, students from the comparison group seemed not to have adequately developed knowledge of logical thinking and reasoning. They did not view the learning of LP from a broader perspective beyond passing national examinations at Uganda Certificate of Education.

The results of this study point to important issues to the educational stakeholders in cultivating an early positive attitude in mathematics which is aimed at investigating specific topics from primary to secondary school mathematics. This may be a potential strategy for applying different heuristic problem-solving approaches and strategies to significantly improve students’ attitudes and performance. Muis et al. (2015) have stated that epistemological beliefs of learners greatly determine the learning strategies that teachers should apply to stimulate their attitude and performance. In this study, the problem-solving heuristic method supported collaboration and discussions between teachers and amongst students during the learning process. It is likely that students from the experimental group worked collaboratively in their small groups. The students helped and guided each other hence boosting their attitude and performance. As noted by Asempapa, (2022), the teachers’ instructional strategies of considering individual students’ differences may change students’ attitude, thereby providing both academic and social support.

Consequently, due to the application of ALHPS approach in the experimental group, the low performing students greatly gained conceptual understanding and also acquired problem-solving skills. This enhanced students’ learning and attitude towards mathematics and LP in particular. Besides, the problem-solving heuristic approach applied to the experimental group boosted students’ confidence in answering both routine and non-routine problems. Students’ fear in comprehending LP word problems and finally attempting to answer LP questions decreased, and their attitude changed too. Students were actively involved in problem-solving. This gradually built students’ competence and confidence in learning LP and related concepts which significantly fostered students’ positive attitude towards LP.

## 6. Conclusion and Future Research Directions

The purpose of this research was to explore the relationship between students’ attitude towards, and performance in LP mathematics word problems. The findings provide preliminary insights into the fundamental concepts and provide an introduction to LP concepts for advanced mathematics. Students’ attitudes point to issues related to the demographic variables and latent constructs for learning mathematics. Attitude towards mathematics has both significant direct and indirect effect on students’ performance mediated by ALHPS strategies. However, some attitudinal dimensions have only direct effect. Wigfield, and colleagues’ expectancy-value model of achievement motivation (Wigfield, 1994; Wigfield & Eccles, 2000) supports this claim. Accordingly, the theory is based on the premise that success on specific tasks and the values inherent in those tasks is positively correlated with performance, and attitude. Thus, the ATMI-SF constructs combining motivation, enjoyment, confidence, and usefulness, and related latent variables are good mediators to explain students’ success in learning LP and mathematics generally. The educational stakeholders and experts in mathematics education should embrace suitable learning strategies that cultivate a positive attitude towards learning and consequently performance.

## Limitations and future research

As earlier mentioned, studies on attitude towards and performance in mathematics have gradually shifted from general to topic domain-specific studies. Thus, instead of investigating the students’ attitude towards mathematics generally, the current research focused on exploring the influences of attitude towards and performance in mathematics word problems and LP in particular. This study was purely quantitative. To gain more insight, we recommend that future researchers should fill the gap through triangulation. The use of qualitative methods such as interviews and observation may provide more evidence on students’ experiences in learning LP word problems, including students’ emotional experiences and the general latent behavior. This would help enrich the existing body of knowledge on attitude towards and performance in mathematics. The teachers ‘attitude towards the domain specific LP word problems is also a potential area for further investigation aimed at improving the instructional strategies. To achieve this, teachers’ content knowledge and pedagogical content knowledge of both in-service and preservice teachers should be investigated. Also, the teachers’ professional development programs should emphasize content knowledge and pedagogical content knowledge. Teachers coming together to share learning experiences and strategies, may help to improve students’ attitude and the learning of presumably “difficult topics” including LP and mathematics generally. Indeed, teachers need routine professional development support to successfully implement the stated learning activities. Another potential area for further research is the relationship between students’ and teachers’ demographic factors mediated by active learning heuristic problemsolving strategies, on students’ performance and attitude towards LP and mathematics generally.

## Acknowledgment

This research is part of the Ph.D. Thesis that investigated the effect of the heuristic problemsolving approach on students’ achievement and attitude towards LP in Ugandan secondary schools. We appreciate useful information provided by the students and teachers in the study sample, which helped us, write this research article. The views expressed herein are those of the authors and not necessarily those of our funders. This is because our funders were not involved in identifying the suitable study design, methods of data collection and analysis, publication decision, or manuscript preparation.

## Author Contribution Statement

Robert Wakhata conceived the research topic, developed methodology, got involved in piloting research tools was involved in investigation, data collection, analysis and Wrote the manuscript. Dr. Védaste Mutarutinya and Dr. Sudi Balimuttajjo did the supervisory role by ensuring that all tools for data collection were validated, reviewed the manuscript and edited and improved subsequent drafts until the manuscript was submitted.

## Data Availability Statement

Data will be made available on request.

## Additional Information

No additional information is available for this paper.

## Appendix 1

### Q1

A factory makes two kinds of bottle tops “Coca-cola” tops and “Pepsi cola tops”. The same equipment can be used to make either. In making Coca-cola tops, one man can supervise 10machines and this batch will give a profit of pounds sterling (£) 50 per week. Pepsi cola tops yield a profit of (£) 250 a week, using 25 machines and 8 men. There are 200 machines and 40 men available. By taking ***x*** batches of Coca-cola tops and ***y*** batches of Pepsi cola tops; write down inequalities for the:

(i) number of machines used (ii) number of men employed (iii) expression for profit, P.

Use these inequalities to draw a suitable graph showing the region which satisfies them. From your graph, determine the numbers of Coca-cola tops and Pepsi cola tops which should be made to obtain the maximum profit. Hence find the maximum profit.

### Q2

A wildlife club in a certain school wishes to go for an excursion to a national park. The club has hired a mini-bus and a bus to take students. Each trip for the bus is Shs.50.000 and that of a mini-bus is Shs.30.000. The bus has a capacity of 54 students and the minibus, 18 students. The maximum number of students allowed to go for the excursion is 216. The number of trips the bus makes do not have to exceed those made by the mini-bus. The club has mobilized as much as Shs.300,000 for transportation of the students. If ***x*** and ***y*** represent the number of trips made by the bus and mini-bus respectively,

a. write down five inequalities representing the above information.
b. plot these inequalities on the same axes.
c. by shading, unwanted regions show the region satisfying the above inequalities.
d. list the possible number of trips each vehicle can make.
e. state the greatest number of students who went for the excursion.

### Q3

A private car park is designed in such a way that it can accommodate ***x*** pickups and ***y*** minibuses at any given time. Each pickup is allowed 15m^2^ of space and each miminibus5m^2^ of space. There is only 400m^2^ of space available for parking. Not more than 35 vehicles are allowed in the park at a time. Both types of vehicles are allowed in the park. But at most 10mini-buses are allowed at that time.

a. i. Write down all the inequalities to represent the above information.
ii. On the same axes plot the graphs to represent the above inequalities in (i) hence shade out unwanted regions.
b. If the parking charges for pick up are Shs.500 and that for a minibus is Shs.800 per day, find how many vehicles of each type should be parked to obtain maximum income, hence find the daily maximum income.

### Q4

A farmer plans to plant an 18hectare field with carrots and potatoes. The farmer’s estimates for the project are shown in the table below.

|  | **Carrots** | **Potatoes** |
| --- | --- | --- |
| Harvesting cost @ ha | Shs.95, 000 | Shs.60,000 |
| Number of working hours | 12days | 4days |
| Expected profit per ha | Shs.228, 000 | Shs.157.000 |

The farmer has shs.1,140,000 only to invest in the project. The total number of working days is 120. By letting ***x*** represent the number of hectares to be planted with carrots and ***y*** the number of hectares to be planted with potatoes

a. write down inequalities for the (i) cost of the project (ii) working days (iii) number of hectares used in the project (iv) the possibility that the field will at least be used for planting either carrots or potatoes.
b. Write down an expression for the profit **P** in terms of ***x*** and ***y***
c. i. On the same axes, plot graphs of the inequalities in (a) and (b) above by shading out the unwanted regions.
ii. Use your graph to determine how the farmer should use the field to maximize profit. Hence, find the farmer’s maximum profit.

### Q5

At a graduation party, the guests are served beer and soda. At least twice as many crates of beer as crates of soda are needed. A crate of beer contains 25 bottles and a crate of soda contains 24 bottles. More than 200 bottles of beer and soda are needed. A maximum of Shs.500, 000 may be spent on beer and soda. Assume a crate of beer costs shs.40, 000 and that of soda costcost.15, 000.

a. i. form inequalities to represent the above information.
ii. represent the above inequalities on the same axes.
iii. by shading the unwanted regions, represent the region satisfying the inequalities in (a) (i) above.
b. From your graph, find the number of crates of beer and soda that should be bought if the cost is to be as low as possible. Find the amount that was paid for those crates of beer and soda.

### Q6

A bicycle factory assembles two types of bicycles, Road master and Hero on different assembly lines. An assembly line for the road masters occupies an area of 60m^2^ of the floor space. The floor space available for all the assembly lines is 420m^2^. The assembly line of the road master needs 10 men to operate it and that of the hero needs 16 men to operate it. The assembly lines need a maximum of 120 men to operate them.

a. If ***x*** and ***y*** represent the number of assembly lines for road master and hero respectively.
  i. form four inequalities to represent the given information.
  ii. draw graphs on the same axes to represent the inequalities in (a) (i) above. Shade the unwanted regions.

### Q7

A private car park is designed in such a way that it can accommodate ***x*** pickups and ***y*** minibuses at any given time. Each pickup is allowed 15m^2^ of space and each miminibus5m^2^ of space. There is only 400m^2^ of space available for parking. Not more than 35 vehicles are allowed in the park at a time. Both types of vehicles are allowed in the park. But at most 10mini-buses are allowed at a time.

a. i. Write down all the inequalities to represent the above information.
ii. On the same axes plot the graphs to represent the above inequalities in (i) hence shade out unwanted regions.
b. If the parking charges for pick up are Shs.500 and that for a minibus is Shs.800 per day, find how many vehicles of each type should be parked to obtain maximum income, hence find the daily maximum income.

## Appendix 2

ATMI-SF Questionnaire for Students’ Attitude towards Linear Programming Word Problems

| No. | Item | SD | D | N | A | SA |
| --- | --- | --- | --- | --- | --- | --- |
| 1. | Linear programming word problems are a very worthwhile and necessary topic in mathematics. |  |  |  |  |  |
| 2. | Linear programming is one of the most important topics in mathematics for students to study. |  |  |  |  |  |
| 3. | Linear programming lessons are very helpful no matter what I decide to study in the future. |  |  |  |  |  |
| 4. | Studying linear programming is important in everyday life. |  |  |  |  |  |
| 5. | Learning linear programming makes me feel nervous. |  |  |  |  |  |
| 6. | I am always under a terrible strain in a mathematics class involving the learning of linear programming word problems. |  |  |  |  |  |
| 7. | I am always confused in my mathematics class whenever we are learning linear programming word problems. |  |  |  |  |  |
| 8. | I feel a sense of insecurity and nervousness when solving a linear programming word problem. |  |  |  |  |  |
| 9. | I have usually enjoyed studying and solving linear programming and related word problems in school. |  |  |  |  |  |
| 10. | Linear programming word problems is one of the interesting topics in mathematics. |  |  |  |  |  |
| 11. | I am always happy, and like learning and solving new tasks in linear programming word problems. |  |  |  |  |  |
| 12. | I am willing to solve and take more than the required amount of linear programming word problems. |  |  |  |  |  |
| 13. | The challenge of learning linear programming word problems appeals to me. |  |  |  |  |  |
| 14. | I plan and am confident to take as many linear programming word problems in advanced mathematics. |  |  |  |  |  |
| 15. | Questions on linear programming should not be made compulsory during national examinations. Please select Agree for this question. |  |  |  |  |  |

## Appendix 3

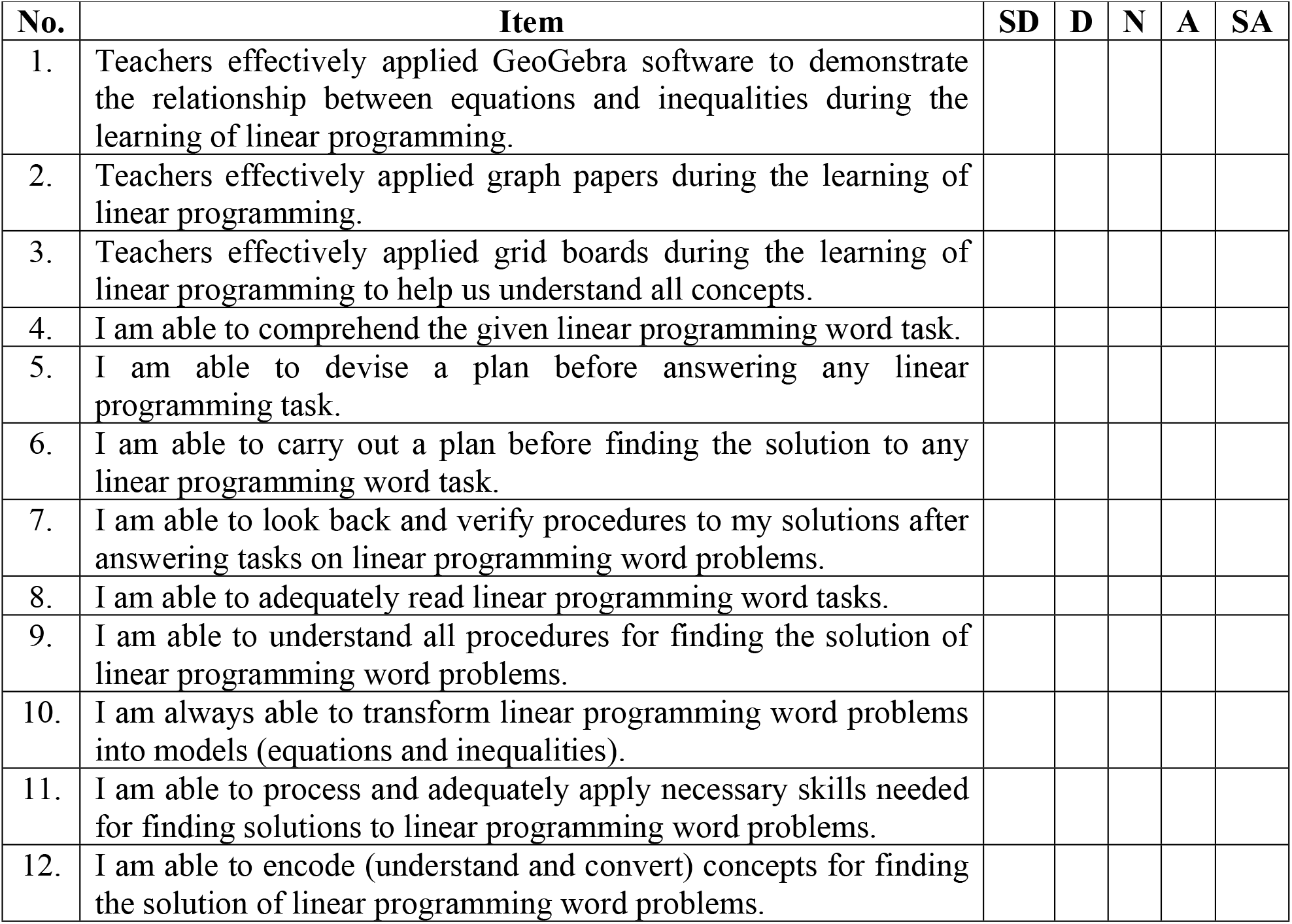
Questionnaire for Students on Application of Active Learning Heuristic Problem-Solving.

Appendix 3 Questionnaire for Students on Application of Active Learning Heuristic Problem-Solving
| No. | Item | SD | D | N | A | SA |
| --- | --- | --- | --- | --- | --- | --- |
| 1. | Teachers effectively applied GeoGebra software to demonstrate the relationship between equations and inequalities during the learning of linear programming. |  |  |  |  |  |
| 2. | Teachers effectively applied graph papers during the learning of linear programming. |  |  |  |  |  |
| 3. | Teachers effectively applied grid boards during the learning of linear programming to help us understand all concepts. |  |  |  |  |  |
| 4. | I am able to comprehend the given linear programming word task. |  |  |  |  |  |
| 5. | I am able to devise a plan before answering any linear programming task. |  |  |  |  |  |
| 6. | I am able to carry out a plan before finding the solution to any linear programming word task. |  |  |  |  |  |
| 7. | I am able to look back and verify procedures to my solutions after answering tasks on linear programming word problems. |  |  |  |  |  |
| 8. | I am able to adequately read linear programming word tasks. |  |  |  |  |  |
| 9. | I am able to understand all procedures for finding the solution of linear programming word problems. |  |  |  |  |  |
| 10. | I am always able to transform linear programming word problems into models (equations and inequalities). |  |  |  |  |  |
| 11. | I am able to process and adequately apply necessary skills needed for finding solutions to linear programming word problems. |  |  |  |  |  |
| 12. | I am able to encode (understand and convert) concepts for finding the solution of linear programming word problems. |  |  |  |  |  |

